# Probing Biomass Precursor Synthesis as a Key Factor in Microbial Adaptation to Unadapted Carbon Sources

**DOI:** 10.1101/2024.10.23.619944

**Authors:** Jingyi Cai, Jiayu Liu, Fan Wei, Wenjun Wu, Wenqi Xu, Zhitao Mao, Qianqian Yuan, Hongwu Ma

**Affiliations:** Biodesign Center, Key Laboratory of Engineering Biology for Low-Carbon Manufacturing, Tianjin Institute of Industrial Biotechnology, Chinese Academy of Sciences, Tianjin 300308, China; National Center of Technology Innovation for Synthetic Biology, Tianjin 300308, China; Tianjin University of Science & Technology, Tianjin 300457, China; University of Chinese Academy of Sciences, Beijing,101408, China

**Keywords:** metabolic network, renewable carbon source, growth coupling, engineering strategies

## Abstract

Industrial microorganisms often face challenges in utilizing renewable substrates such as methanol, formate, and xylose. We present findings that the proportion of biomass precursors that must be synthesized from unadapted carbon sources is a critical determinant of the evolutionary driving force and minimal substrate requirements, using a new computational framework, AdaptUC. We predict metabolic engineering strategies for Adaptive Laboratory Evolution (ALE). These strategies enable microorganisms to co-utilize an adapted co-substrate and an unadapted carbon source or, in some cases, rely exclusively on the unadapted source. AdaptUC was validated through experimental records and literature, confirming its effectiveness in identifying gene knockout strategies. Case studies in *Escherichia coli* and *Corynebacterium glutamicum* highlight superior strategies with higher driving forces and reduced substrate requirements. This method has the potential to transform industrial biosynthesis by enabling more efficient use of renewable carbon sources.

**Graphic Abstract:** 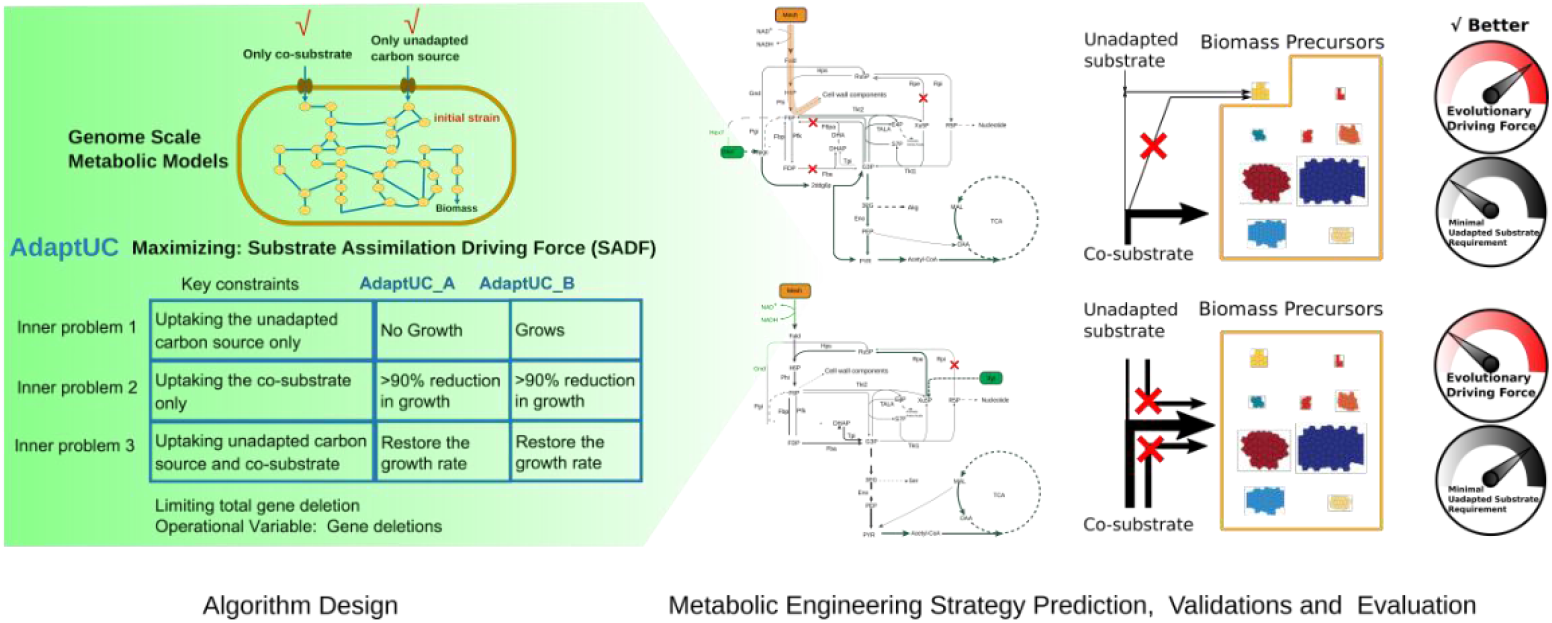

## INTRODUCTION

Industrial microorganisms are essential for sustainable biosynthesis processes, but they often struggle to utilize renewable substrates such as methanol, formate, and xylose. These substrates—derived from non-food materials or industrial waste like methanol (oxidized from methane or synthesized from carbon dioxide^1^), xylose (from agricultural waste), glycerol (a biodiesel by-product), formate (electrochemically synthesized from carbon dioxide ^2^) hold great promise for industrial biosynthesis. However, many industrial strains cannot naturally assimilate these unadapted carbon sources due to factors such as the absence of assimilation pathways, redox imbalance^3^, unfavorable enzyme kinetics ^4^, inadequate enzyme expression levels, or regulatory mechanisms ^5^. Ultimately, this lack of adaptation is likely because these substrates were not present in the microorganisms’ natural environments during evolution ^6^. An immediate solution is to introduce heterologous genes encoding the necessary pathways into the host cell, which has been effective in enabling *C. glutamicum* to assimilate xylose ^7^, and *E. coli* to grow on serine ^8^. Nevertheless, a complete assimilation pathway may not suffice for growth on an unadapted carbon source. For example, the toxicity of intermediate formaldehyde still prevents unevolved *E. coli* from growing when a heterologous methanol utilization pathway is inserted ^9,10^. Similarly, *E. coli* cannot utilize glycerol under anaerobic conditions due to redox imbalance and thermodynamic barriers ^3^ due to redox imbalance and thermodynamic barrier, *E*.*coli* grows poorly on formate as a sole carbon source without ALE^11,12^.

To overcome these challenges, it is crucial to understand the factors determining the evolutionary driving force and minimal substrate requirements for the assimilation of unadapted carbon sources. ALE is a powerful technique that helps cells adapt and fine-tune their cellular machinery for better growth on new substrates. Given the starting strain’s difficulty in assimilating unadapted carbon sources, favored co-substrates are required to support its growth initially. This is followed by gradually changing environmental conditions to force the strain to evolve towards a phenotype with high utilization capacity for the previously unadapted substrate.

A critical question in this context is how the proportion of biomass precursors that must be synthesized from unadapted carbon sources affects the performance of metabolic engineering strategies. This question is significant because if the proportion of biomass precursors from unadapted substrates plays a major role, it could serve as a guiding principle in designing new microbial strains capable of efficiently synthesizing chemicals from previously unadapted renewable substrates. By carefully designing gene knockout strategies that couple cell growth and substrate assimilation, we can increase the evolutionary driving force. This strategy has been successfully demonstrated in creating methylotrophic E. coli that efficiently utilize methanol as the sole carbon source ^9^ and in *C. glutamicum* to achieve co-utilization of methanol and xylose^13^.

Computational tools can significantly enhance the design of starting strains for the ALE process. However, current methods are limited by their reliance on small-scale core metabolic models to avoid combinatorial explosion, making exhaustive searches impractical in genome-scale models ^5^. For example, Keller et al. used the *E. coli* core model to evaluate 33 gene knockout combinations across 11 co-substrates, resulting in over 2 million in silico experiments using Flux Balance Analysis (FBA) ^14,15^. However, the comprehensive genome-scale metabolic model of *E. coli*, iML1515, with 2712 reactions, makes exhaustive searches impractical. Furthermore, no computational work has assessed the minimal requirement of unadapted carbon sources for the initial strain or the substrate assimilation driving force under the designed genotypes.

In this study, we present a new computational framework, AdaptUC, comprising two variants of bi-level mixed-integer programming (MIP) algorithms. AdaptUC is designed to uncover insights into microbial carbon source adaptation by identifying gene knockout strategies that serve as an evolutionary driving force for strains to utilize previously unadapted carbon sources. By transforming all considerations for the design of the initial strain into internal constraints that must be simultaneously satisfied, AdaptUC can identify knockout targets that couple the target substrate with growth in genome-scale metabolic models containing thousands of reactions. Two quantitative metrics are used to evaluate the predicted strategies: the minimal requirement of the unadapted carbon source for the initial strain and the carbon assimilation driving force correlated with the degree of growth coupling.

We demonstrate the effectiveness of AdaptUC through case studies involving important industrial species, *Escherichia coli* and *Corynebacterium glutamicum*, focusing on methanol utilization both in co-utilization with other carbon sources and as the sole carbon source. We validate our method by comparing the prediction results with experimentally effective strategies reported in the literature. Furthermore, we present new strategies with superior potential, characterized by higher evolutionary driving forces and lower initial substrate requirements, found through an in-depth metabolic-level analysis. Our findings reveal that the proportion of biomass precursors that must be synthesized from unadapted carbon sources is a critical determinant of the evolutionary driving force and minimal substrate requirements. By leveraging computational analysis through AdaptUC, we can predict metabolic engineering strategies for ALE, enabling microorganisms to co-utilize an adapted co-substrate and an unadapted carbon source or, in some cases, exclusively rely on the unadapted source.

## RESULTS AND DISCUSSION

### Design of AdaptUC

In this study, we present a computational framework named AdaptUC, designed to predict gene knockout (KO) strategies for developing a starting strain for the ALE process. The goal is to enable the utilization of an unadapted carbon source. As depicted in Figure 1, AdaptUC includes two algorithms: AdaptUC-A and AdaptUC-B. AdaptUC-A, with its concept initially introduced by Gawand et al ^16^, couples strain growth with both the unadapted substrate and a co-substrate, requiring simultaneous supply of both for growth. AdaptUC-B, a new bi-level alogrithm couples strain growth exclusively with the unadapted substrate, enabling growth only when the unadapted substrate is present. AdaptUC-A predicts knockout targets that result in the strain requiring both the unadapted substrate and the adapted co-substrate for growth. This approach allows the strain to gradually adapt to the unadapted substrate during the evolutionary process, although it may still be unable to grow on the previously unadapted substrate as the sole carbon source. In contrast, AdaptUC-B preserves the potential for utilizing the unadapted substrate as the sole carbon source. Both AdaptUC_A and B include three inner problems (design requirements), each with an objective that must satisfy the constraints in the outer level with minimal number of modifications (Figure 1). Each inner problem and its corresponding outer-level constraint represents a requirement for the ALE starting strain, delineating the expectations for the strain at the conclusion of the ALE process.

**Figure 1.**
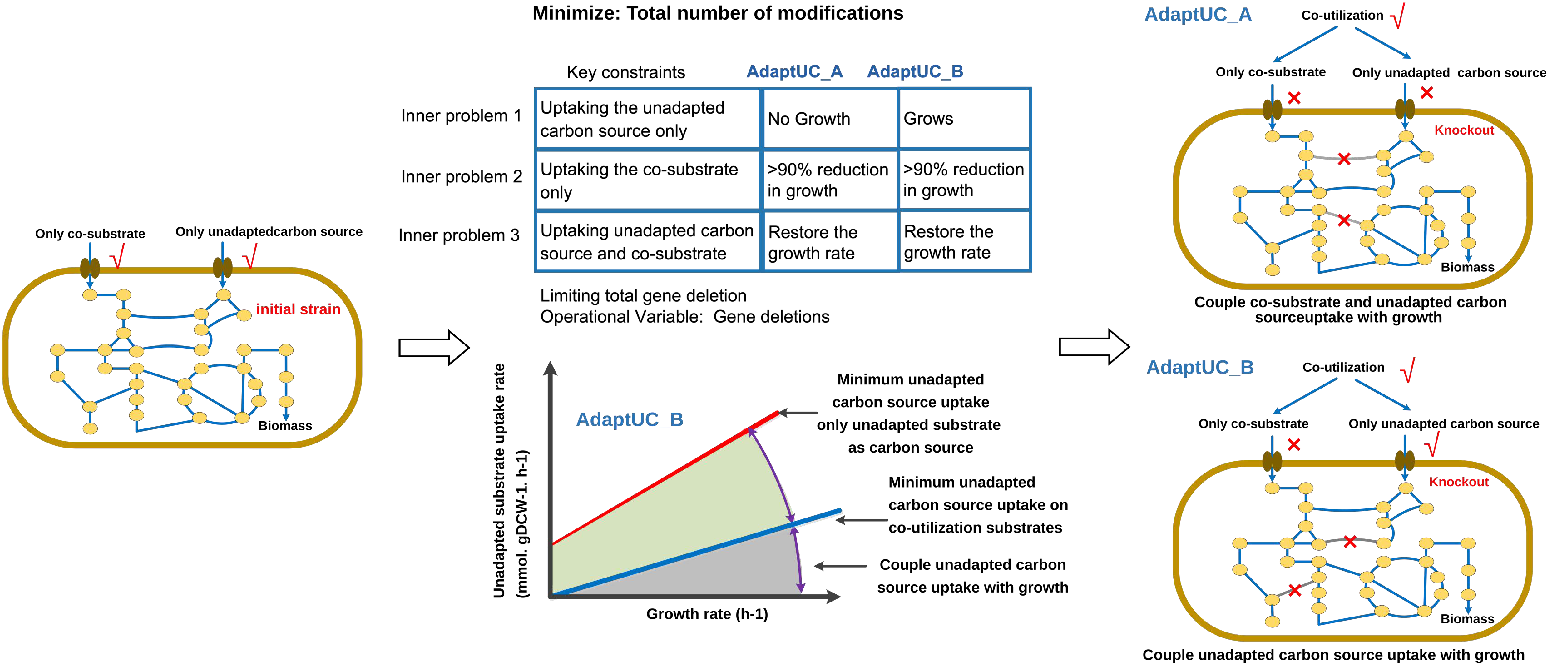
The schematic diagram for the concept of AdaptUC.

The first inner problem signifies the necessity for the strain after ALE to exhibit no growth (AdaptUC-A) or robust growth (AdaptUC-B) on a medium with an unadapted substrate as the sole carbon source (Figure 2). The second requirement dictates that the strain after ALE should not thrive well without the unadapted carbon source, even in the presence of the co-substrate. Combining these two requirements effectively couples the assimilation of the unadapted carbon source with growth, generating a compelling driving force for the cell to uptake the unadapted substrate. In the final inner problem, where the unadapted substrate is available for uptake, as opposed to the second inner problem, the objective is to ensure that the optimal growth of the strain after ALE is not inferior to the growth of the reference strain on the co-substrate. This design requirement guarantees that the addition of conventional substrate to the medium containing the unadapted carbon source can aid the survival and growth of the starting strain in the ALE process. We assume that the gene deletions for the strain after ALE and the ALE starting strain are either similar or that their differences do not significantly impact growth. AdaptUC is specifically designed to construct the ALE starting strain. The algorithm details are provided in the Methods section.

**Figure 2.**
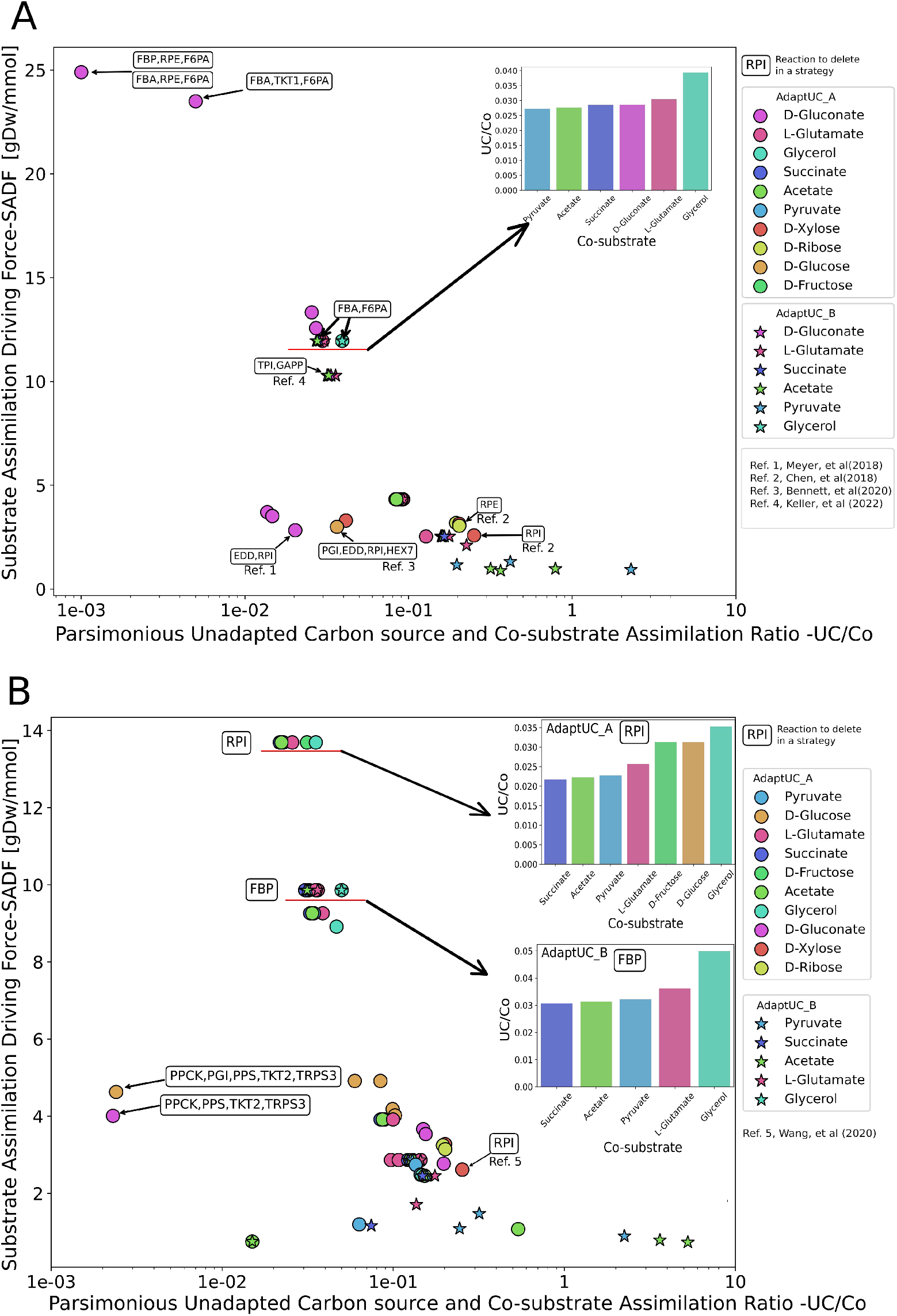
Substrate assimilation driving force of AdaptUC predicted strategies for designing methylotrophic *(A) E. coli* and *(B) C. glutamicum*. Successful strategies reported by literature are annotated, original data in SI Table 8 and SI Table 9

### Designing metabolic engineering strategies for creating methylotrophic strains and literature validation

We applied AdaptUC to predict gene modifications for developing methylotrophic *E. coli* strains. Methanol serves as an unadapted carbon source for *E. coli*, which naturally lacks a methanol assimilation pathway. The mere insertion of a heterogeneous methanol assimilation pathway doesn’t immediately grant *E. coli* the ability to grow on a medium where methanol is the sole carbon source ^9^. This necessitates gene modifications and Adaptive Laboratory Evolution (ALE) experiments to induce *E. coli* to evolve the capacity for methanol assimilation. To facilitate this, we first introduced the methanol assimilation pathway (comprising methanol dehydrogenase, 3-hexulose 6-phosphate synthase, and 6-phospho 3-hexulose isomerase) into the most recent *E. coli* K12 model, *iML1515*. We then used AdaptUC to predict gene modification strategies for constructing the initial strain. To ensure the initial strain’s survival in the initial ALE media, we considered 10 co-substrates (succinate, fructose, glucose, acetate, glutamate, glycerol, glycerate, pyruvate, xylulose, ribose). The number of allowable knockout reactions were limited to no more than 5. This analysis yielded 89 solutions, evaluated based on two factors: the parsimonious Unadapted Carbon source and Co-substrate assimilation ratio (UC/Co, see method) on the x-axis and the average Substrate Assimilation Driving Force (SADF, see method) of methanol on the y-axis (Figure 2A). AdaptUC successfully predicted five experimentally validated strategies for methylotrophic *E. coli* strains (SI, table 1). Four of these are AdaptUC-A strategies, disrupting the RuMP cycle to prevent the formaldehyde acceptor ru5p from being generated by methanol alone, while preventing the co-substrate from entering the EMP or ED pathway without methanol, forcing the co-substrate to be the sole precursor of *ru5p*. The validated RuMP-interrupted strategies include Xylose-cosubstrate-Δ*rpi*^17^, Ribose-cosubstrate-Δ*rpe* ^17^, and glucose-cosubstrate-Δ*pgi*Δ*edd*Δ*rpi* ^18^ and Gluconate-cosubstrate-Δ*edd*Δ*rpi* ^19^. An AdaptUC-B strategy, using pyruvate as the co-substrate and deleting *tpi*, was also validated by literature ^9^. Due to limitation of the stoichiometric model, there are minor differences between literature and the predicted genotype, which are explained in the SI notes.

We predicted strategies with much higher SADF and lower UC/Co than reported strategies. For *E*.*coli*, deletion of *fba, rpe*, and *f6pa* while using gluconate as the co-substrate shows a more than fivefold higher driving force than currently reported AdaptUC-A strategies (Figure 2A) with the same deletions. The selection of co-substrate can significantly affect SADF. For the AdaptUC-B strategies, we predict that the pyruvate-Δ*fba*, Δ*f6pa* strategy has slightly higher SADF and lower UC/Co than the experimentally successful co-substrate-pyruvate-ΔTPI strategy, warranting an experimental testing.

For *C glutamicum*, there are significantly fewer reported cases compared to *E. coli*. Deletion of Ribose-5-phosphate isomerase (*rpi*) and using xylose as a substrate can achieve co-utilization of xylose and methanol, a reported case that has been predicted by AdaptUC. However, similar to the situation in *E. coli*, we found that in the deactivation of *rpi*, xylose can only be converted to ru5p and combine with formaldehyde in a 1:1 ratio, causing the intake of formaldehyde and xylose to be completely coupled. Compared to strategies that decouple methanol from the co-substrate, achieving the same growth rate with this strategy requires more methanol, resulting in a lower SADF. We also predicted that simply changing the co-substrate while keeping the *rpi* deletion can significantly decouple methanol from the co-substrate, improving SADF and lowering UC/Co. Predictions show that using seven different substrates, such as succinate, acetate, and pyruvate, can achieve this decoupling effect, increasing SADF by 5.2 times and reducing UC/Co by 91% (Figure 2B).

*C. glutamicum* requires fewer reaction deletions compared to *E. coli*. This is because *C. glutamicum* lacks the Entner-Doudoroff (ED) pathway and some gluconeogenesis reactions present in *E. coli*. For example, within the AdaptUC-B strategy, the preferred approach is the deletion of *FBP*. This strategy allows for the production of fructose-6-phosphate, which is necessary for cell wall synthesis, without disrupting the exclusive utilization of methanol to produce it. Additionally, there is no need to delete an alternative glyconeogenesis fructose 6-phosphate aldolase (*F6PA*) because it is not present in *C. glutamicum*.

Simplified metabolic flux diagrams have been prepared for four unreported promising strategies for *E. coli* and *C. glutamicum* (Figure 3). The top-right of each sub-figure shows the minimal uptake rates of methanol as a function of growth rate under different co-substrate uptake rates. A positive reciprocal of the slope indicates the degree of growth coupling with methanol uptake rates. The slope may change with increasing growth rates, signifying a switch in metabolic flux patterns. As the slope increases, it indicates that co-substrates are becoming insufficient, and the unadapted carbon source must be utilized to synthesize more biomass precursors. During the transition from growing on both co-substrates and the unadapted substrate to growing solely on the previously unadapted substrate, the flux direction in the lower EMP pathway may change, requiring regulatory adjustments during the ALE process (Figure 3 B/C and E/F).

**Figure 3.**
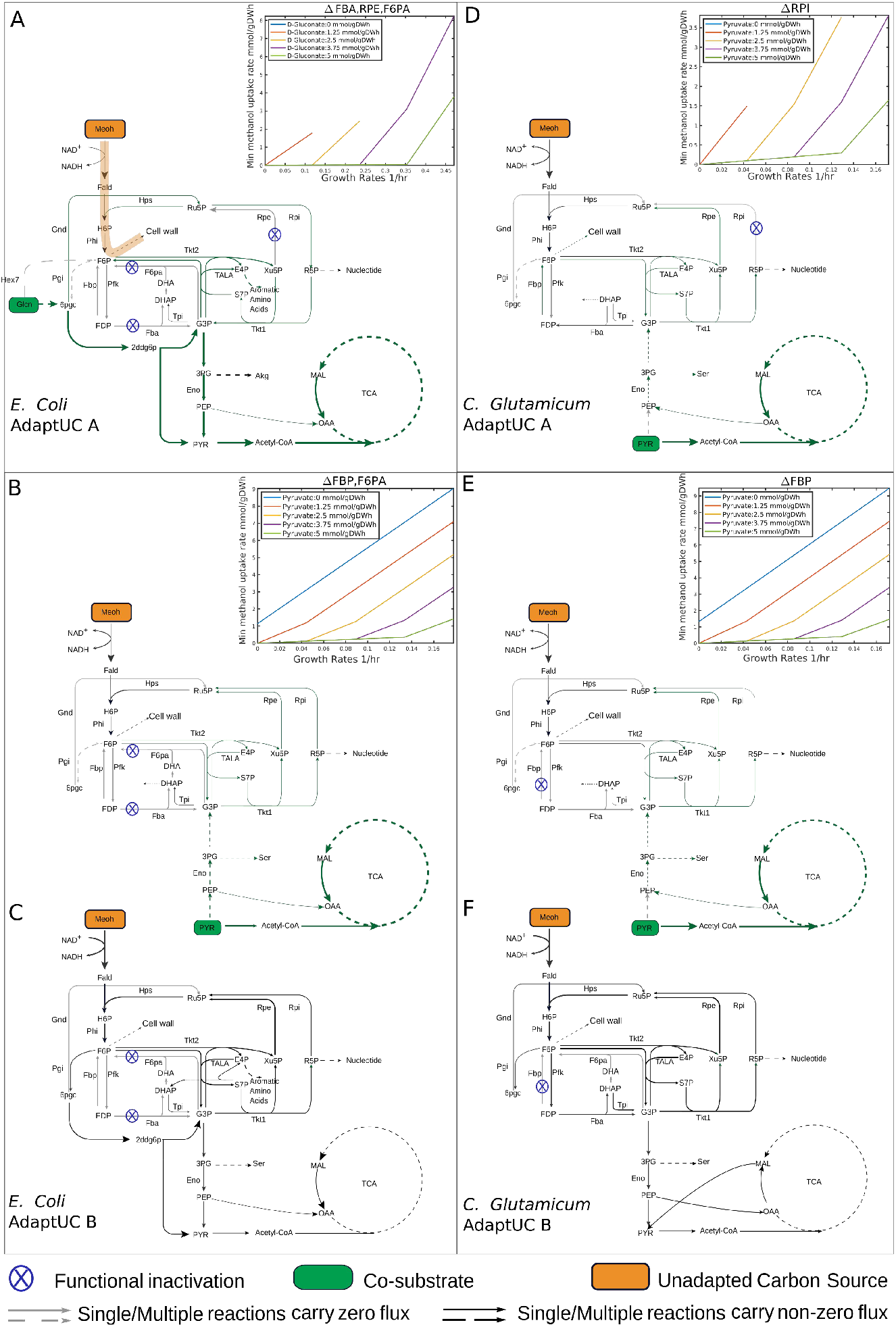
Selected promising strategies for interrupted (A, D) and uninterrupted (B, C, E, F) RuMP cycle corresponding to AdaptUC-A and AdaptUC-B strategies, respectively. For AdaptUC-B strategies, panels B and E show the flux diagrams before evolution, where the co-substrate and unadapted carbon source are utilized simultaneously. Panels C and F show the flux diagrams after evolution, where methanol (the unadapted carbon source) is used as the sole carbon source. The smaller plots in the top-right of each sub-figure depict the minimal uptake rates of methanol as they change with growth rate under different co-substrate uptake rates.

### A workflow combining AdaptUC-A and AdaptUC-B

We observed that for both *E. coli* and *C. glutamicum*, AdaptUC-A strategies can yield higher SADF and lower UC/Co ratios compared to AdaptUC-B strategies (Figure 2). There are overlaps in targets within AdaptUC-A and AdaptUC-B strategies. For example, one of the best AdaptUC-A strategies for *E. coli* with gluconate as a co-substrate involves Δ*fba*Δ*rpe*Δ*f6pa*, achieving an SADF of 24.9 and a UC/Co of 0.001. In contrast, one of the best AdaptUC-B strategies for *E. coli* using pyruvate as a co-substrate involves Δ*fba*Δ*f6pa*, with a lower SADF of 11.9 and a higher UC/Co of 0.027. This indicates that if AdaptUC-B strategies fail, we can switch to an AdaptUC-A strategy that requires less unadapted carbon source (lower UC/Co) and has a higher evolutionary driving force (SADF). Through this process, the strain can adapt to the unadapted substrate more easily. If the AdaptUC-A strategy is successful in achieving a strain that grows solely on an unadapted carbon source, minimal adjustments can be made by comparing both AdaptUC-A and AdaptUC-B strategies. In this example, rpe should be reintroduced to the strain, and the co-substrate should be gradually changed to pyruvate, followed by evolving the Δ*fba*Δ*f6pa* strain, making the strain’s growth dependent on the unadapted substrate until the co-substrate is finally reduced to zero. We have summarized the workflow in Figure 4 to maximize the success of producing strains capable of growing on a medium with an unadapted substrate as the sole carbon source.

**Figure 4.**
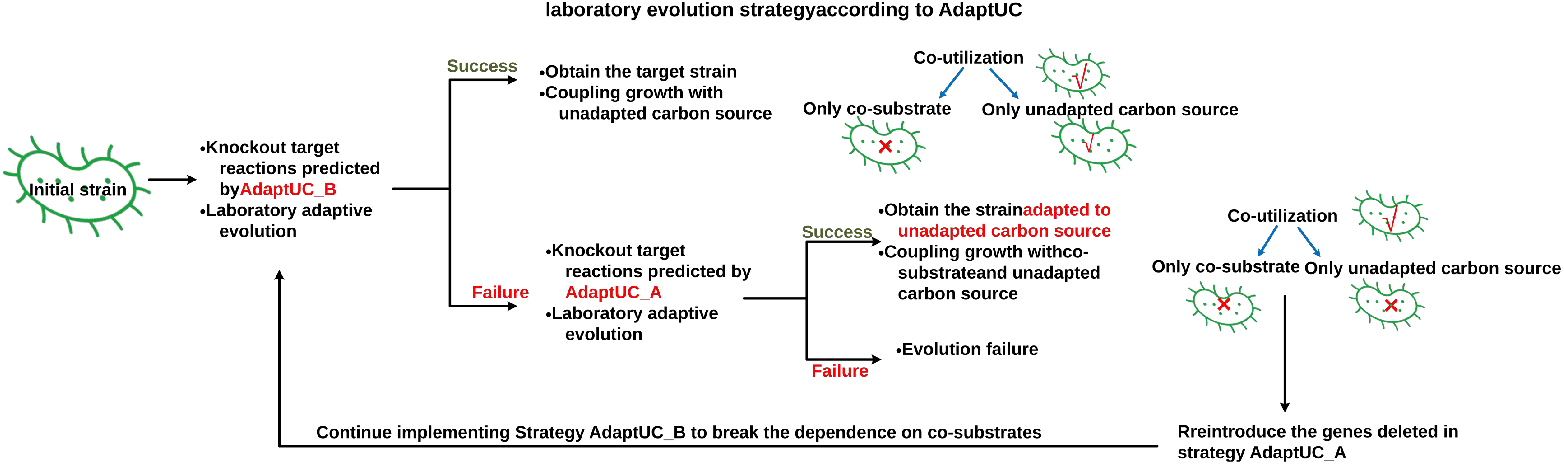
The proposed workflow combines AdaptUC-A and AdaptUC-B to maximize the success of producing strains capable of growing on a medium with an unadapted substrate as the sole carbon source.It is advisable to initially attempt the AdaptUC-B strategy, since it preserve the potential of the strain growing solely on the unadapted substrate.

### The degree of growth coupling of unadapted carbon source determine the evaluation metrics of strategies

A desirable strategy is characterized by a high Substrate Assimilation Driving Force (SADF) and a low UC/Co ratio. From Figure 2, it can be observed that the strategies are not evenly distributed in the SADF and UC/Co panel but are clustered together. the SADF and UC/Co values of strategies within different cluster vary significantly. By analyzing metabolic flux pathways, we discovered that the substantial differences in SADF and UC/Co values among strategies in different cluster are due to the varying degrees of coupling between the unadapted substrate and cell growth. Strategies clustered together in Figure 2 exhibit similar degrees of growth coupling, meaning that the same or similar biomass precursors must be synthesized using the unadapted substrate. Both AdaptUC-A and AdaptUC-B ensure that not all biomass components can be synthesized solely from the co-substrate to create a certain level of substrate utilization driving force. Consequently, the growth coupling degree of the unadapted substrate in all predicted strategies ranges from 0% to 100%. A low degree of growth coupling indicates that only a small portion of the biomass requires the unadapted substrate for synthesis. In such cases, the UC/Co ratio is low, suggesting that the initial strain can easily survive on media containing the unadapted substrate. Simultaneously, most of the biomass is synthesized from the co-substrate, which the cell can assimilate in large amounts. This means that even a small intake of the unadapted substrate can significantly boost biomass production, resulting in a high SADF. We classify strategies into three categories based on the degree of growth coupling with the unadapted substrate: minimal coupling, intermediate coupling, and full coupling. Full coupling means that the synthesis of all biomass precursors depends entirely on the unadapted substrate (Figure 5).

**Figure 5.**
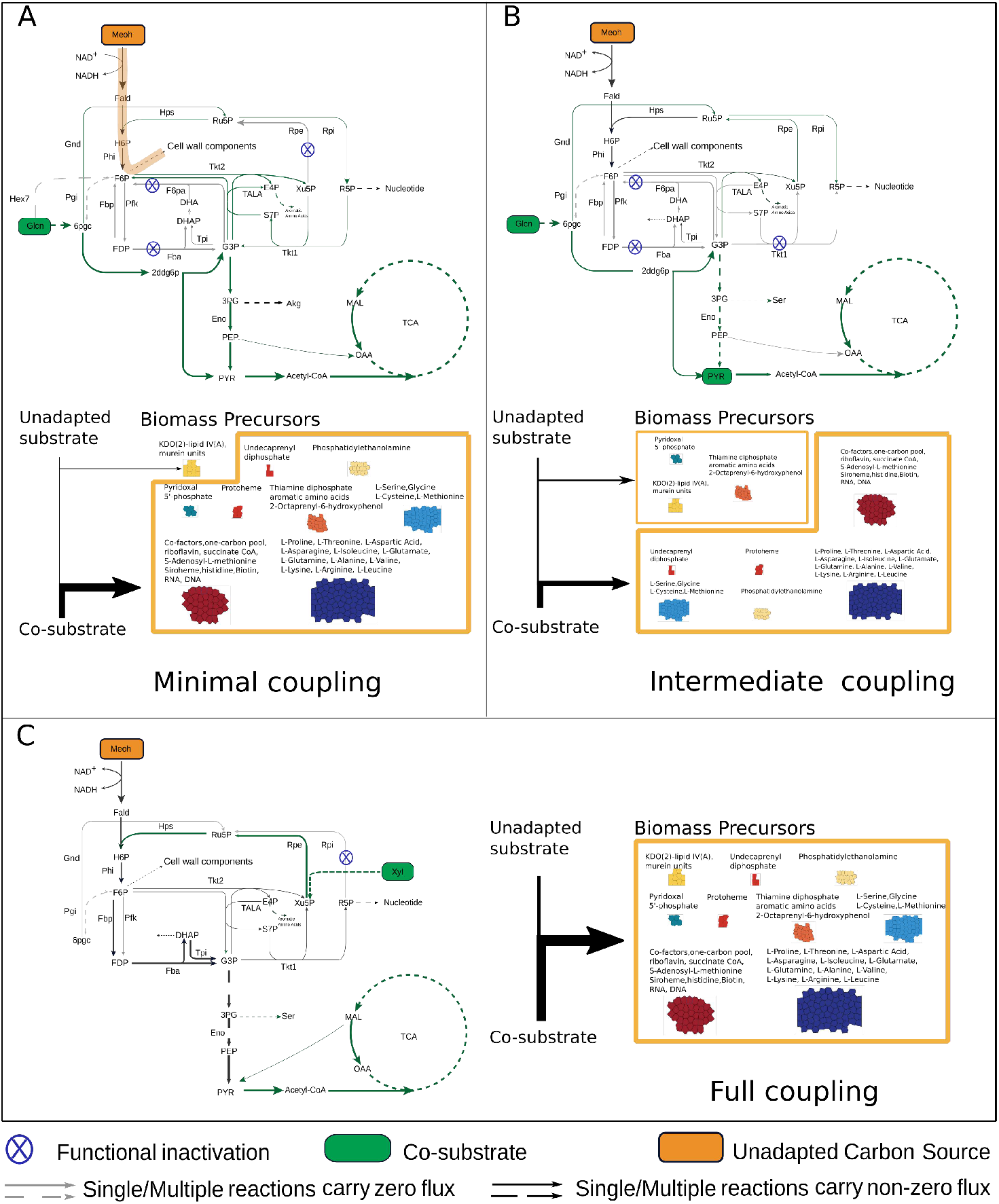
Mode of unadapted carbon source coupling to biomass, A, minimal coupling, only mimimal portion of biomass must be synthesized with an unadapted carbon source. B, intermediate coupling, multiple category but not all of biomass components must be synthesized with an unadapted carbon source. C, full coupling, no biomass components can be produced without the unadapted carbon source.

Specifically, the minimal coupling case for E. coli involves using gluconate as a substrate and knocking out *fba* (fructose-bisphosphate aldolase), *f6pa* (fructose 6-phosphate aldolase), and *rpe* (ribulose 5-phosphate 3-epimerase) (Figure 5A). In this scenario, due to the deletion of *fba* and *f6pa*, gluconate-derived glyceraldehyde 3-phosphate cannot produce D-fructose 6-phosphate. However, D-fructose 6-phosphate is an irreplaceable precursor for KDO(2)-lipid IV(A) and murein units, which contribute respectively to the peptidoglycan layer and lipopolysaccharide (LPS) structure of the cell wall. KDO(2)-lipid IV(A) and murein units represent only 5.48 wt% of the biomass. Therefore, D-fructose 6-phosphate synthesis must occur through formaldehyde (derived from the unadapted substrate methanol) reacting in a 1:1 molar ratio with D-ribulose 5-phosphate. The latter is entirely derived from gluconate, but only accounts for 0.63% of the gluconate pathway. The remaining 92.3% of the carbon flux channels through the Entner-Doudoroff pathway towards glyceraldehyde 3-phosphate and downstream glycolysis and the TCA cycle, while 7.1% converts to D-ribose 5-phosphate, which later channels into nucleic acid synthesis. The synthesis of KDO(2)-lipid IV(A) and murein units, which make up only 5.48 wt% of the biomass, requires methanol (Figure 6A and B). Due to the low proportion of these biomass precursors and the fact that 83% of the carbon comes from gluconate, this minimal coupling strategy results in a very low UC/Co (0.001) and a high SADF (24.9 gDW/mmol).

**Figure 6.**
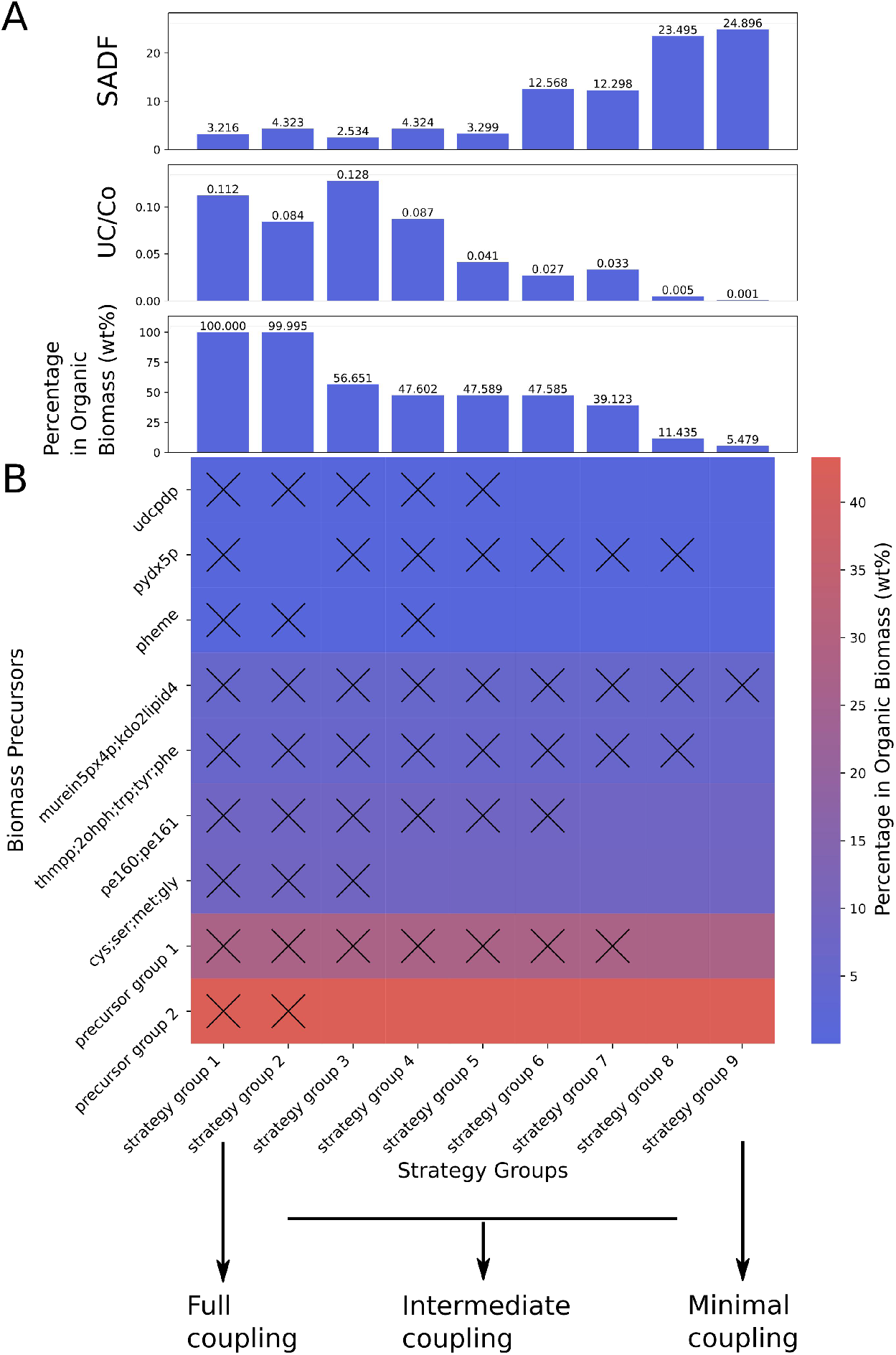
Biomass precursors producibility from co-substrate for AdaptUC-A predicted strategies. A. the SADF, UC/Co, and percentage of biomass precursors that must be synthesized with unadapted precursors. B. the color in the heat map indicate the mass percentage of precursor groups in total organic cellular biomass of *E. coli*, the cross mark indicate the corresponding biomass precursor group can’t be synthesized by co-substrate alone (or must be synthesized by the unadapted carbon source) under the corresponding strategy group in x axis. Precursor group 1: biotin, riboflavin, succoa, amet, thf, mlthf, 10fthf, fad, sheme, nadp, coa, nad, dttp, dctp, datp, dgtp, histidine, ctp, utp, gtp. Precursor group 2: pro, thr, asp, asn, ile, glu, gln, ala, val, lys, arg, leu. Strategy Groups 9 (minimal coupling): glcn: FBA, RPE, F6PA; glcn: FBP, RPE, F6PA, Strategy Groups 1 (full coupling): xyl:TKT1;xyl:RPI et al totally 16 strategies. The full content of each strategy group can be found in SI Table 6-7.

In contrast, a full coupling scenario has a much lower driving force and a much higher UC/Co ratio. For example, in the case of deleting *RPI* while using D-xylose as the co-substrate (Figure 5C), D-xylose cannot synthesize biomass precursors independently but channels to D-ribulose 5-phosphate, which can only combine with formaldehyde to form hexulose 6-phosphate in a 1:1 molar ratio, This results in no biomass precursor being synthesized without methanol (Figure 6AB the most left column). Most predicted strategies are intermediate coupling to growth (Figure 5 B, and Figure 6A/B the middle columns). Generally, SADF increases and UC/Co decreases with the decrease of the degree of the unadapted carbon coupling with growth. Strategies with low growth coupling are advantageous (Figure 6A).

## CONCLUSION

AdaptUC streamlines the design of starting strains for the ALE process, predicting gene knockout strategies that serve as an evolutionary driving force for strains to utilize previously unadapted carbon sources. By focusing on methanol utilization, both in co-utilization and sole utilization scenarios, AdaptUC successfully predicts experimentally validated strategies and identifies superior strategies with higher evolutionary driving forces and lower initial methanol requirements.

Our findings highlight that the proportion of biomass precursors synthesized from unadapted carbon sources plays a critical role in determining the evolutionary driving force and minimal substrate requirements. By leveraging computational analysis through AdaptUC, we can predict metabolic engineering strategies for ALE, enabling microorganisms to co-utilize an adapted co-substrate and an unadapted carbon source or, in some cases, exclusively rely on the unadapted source.

This method has the potential to transform industrial biosynthesis by enabling more efficient use of renewable carbon sources. It may be possible to establish a strain library by leveraging high-throughput automated biofoundry systems in combination with strain construction scheduling optimization algorithms ^20^, This approach would allow for the identification of the optimal strains to support synthetic biology research focused on product synthesis.

## METHOD

### Development of AdaptUC

The two components of AdaptUC framework, AdaptUC-A and AdaptUC-B, each encompasses three requirements that must be satisfied simultaneously. The first design requirement is that the strain after gene modification has the potential to evolve to a phenotype that can grow on the untapped carbon source. The growth rate is simulated by flux balance analysis.

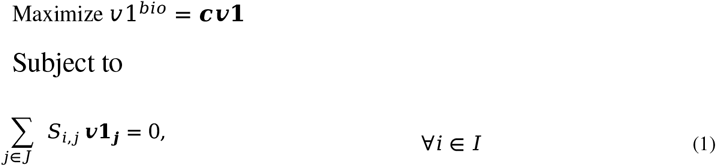

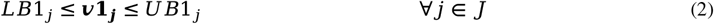

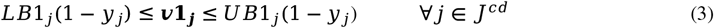

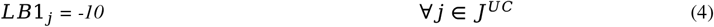

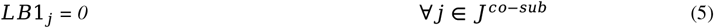

Where *S*_*i,j*,_ denotes the stoichiometric coefficient matrix of metabolite i in reaction j, ***v*1**_***j***_ is the flux variable of reaction j for the first design requirement, *LB*1_***j***_ and *UB*1_***j***_ are lower and upper bounds for ***v*1**_***j***_, these bounds can be used to setup reactions’ reversibility based on thermodynamics. Where the binary variable *y*_*j*_ indicates whether the reaction j is deactivated (*y*_*j*_ =1) or not (*y*_*j*_ =0).

The outer level constraint on the first inner problem is that, for AdaptUC B variant, it must still can grow on the unadapted substrate after deletion, the reduction of growth due to gene deletion must not exceed 50% :

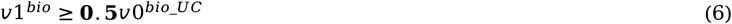

Where *v*0^*bio_UC*^ is the growth rate of reference strain on the untapped carbon source that can be calculated by solving the following FBA problem:

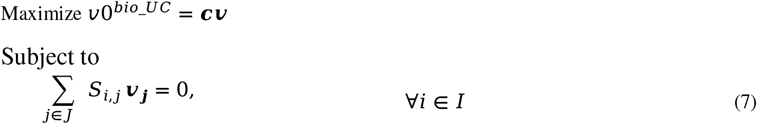

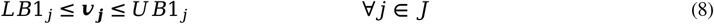

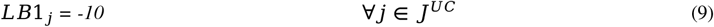

Where *J*^*UC*^ is and are uptake reaction(s) for unadapted substrate and **c** is the objective coefficient vector.

For AdaptUC A variant, the deletion cause the strain being not able to grow on the unadapted substrate, we set the growth reduction greater than 90% to achieve this constraint.

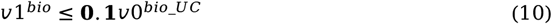

The second requirement is that after deletion, the strain suffer no growth or severely impaired growth on conventional co-substrate, forcing the strain to utilize C1-substrate whenever it’s available. Considering reduction of 90% in growth fulfil this requirement, then this requirement is implemented in the following constraint:

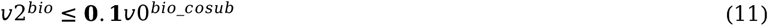

The calculation for growth rate of the reference strain on co-substrate is similar to *v*0^*bio_UC*^, the difference is to setting the bound for co-substrate exchange reaction instead. The growth rate *v*2^*bio*^ is calculated by the following inner problem:

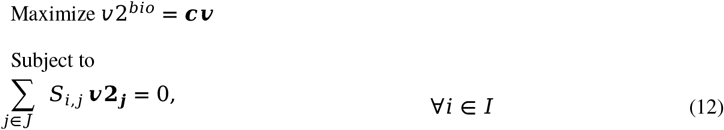

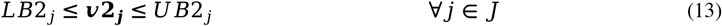

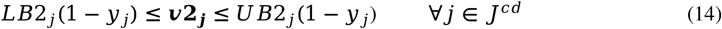

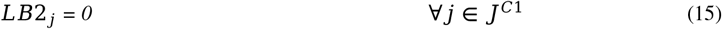

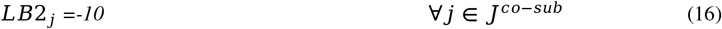

The third requirement is that addition of unadapted substrate to the co-substrate-only medium can recover the growth rate to the growth before gene deletion. This requirement is implemented by the following outer-level constraint:

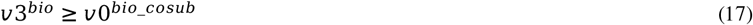

Where the growth rate *v*3^*bio*^ is calculated by the third inner problem:

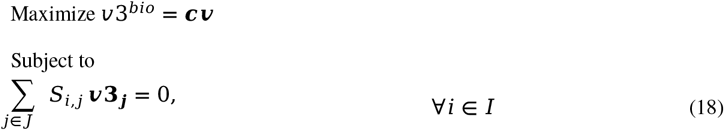

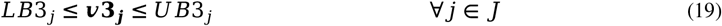

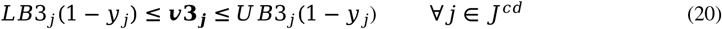

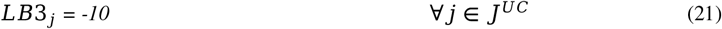

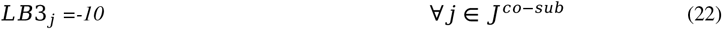

The minimal uptake rate for the unadapted carbon source to recover growth after gene deletion, *v*^*UC*^ can be obtained by the following inner problem:

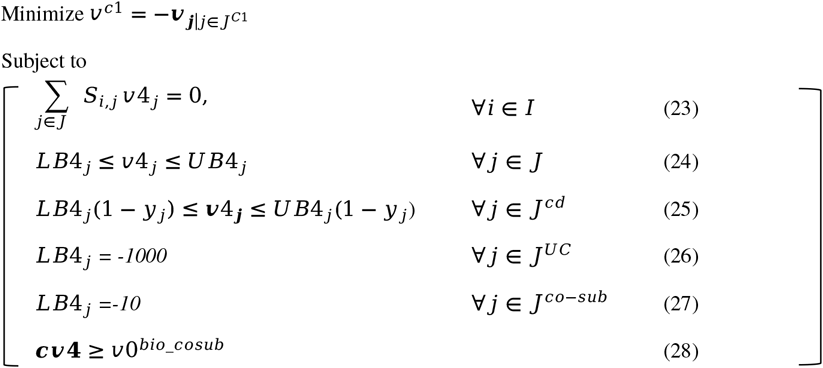

The total number of reaction deletion must be constrained to accelerate the solving process:

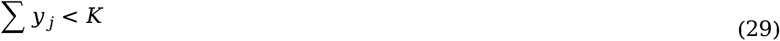

In summary, the algorithm AdaptUC is then formulated as below:

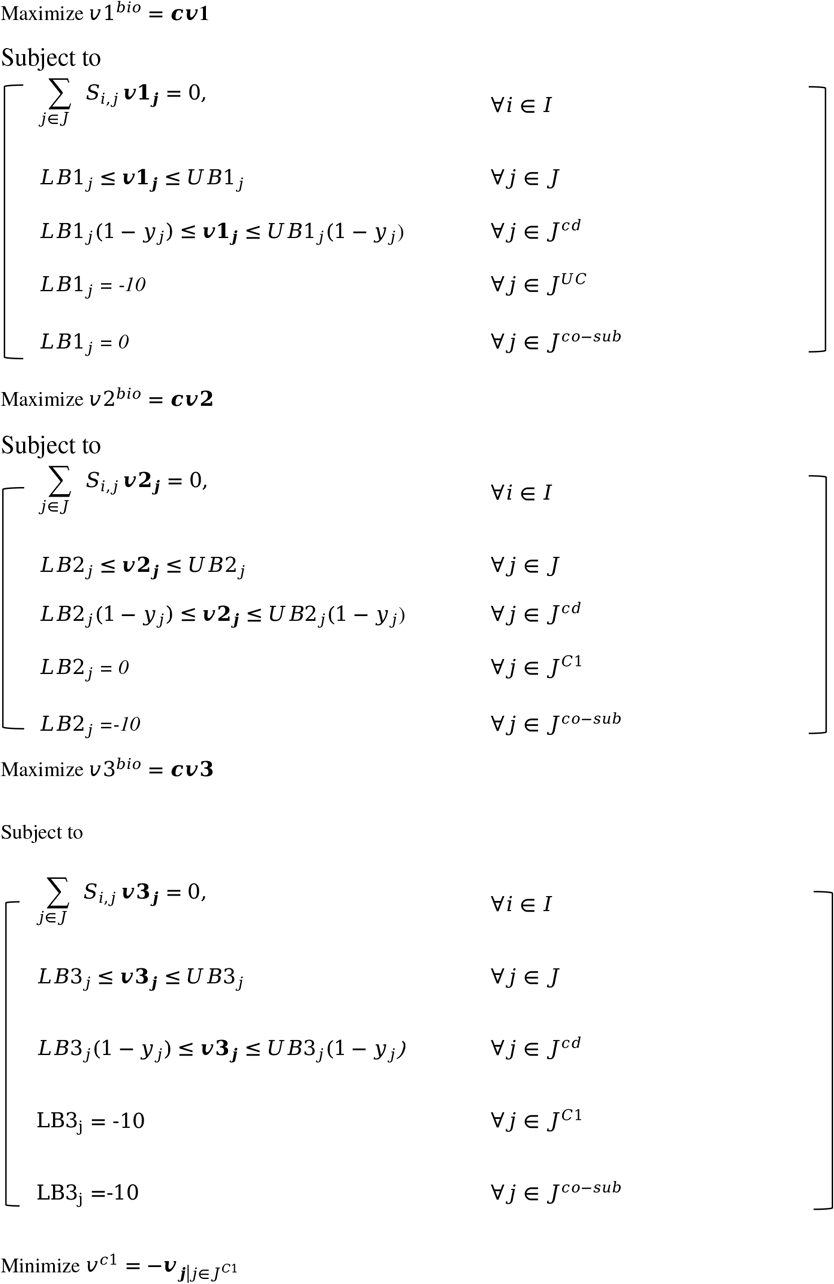

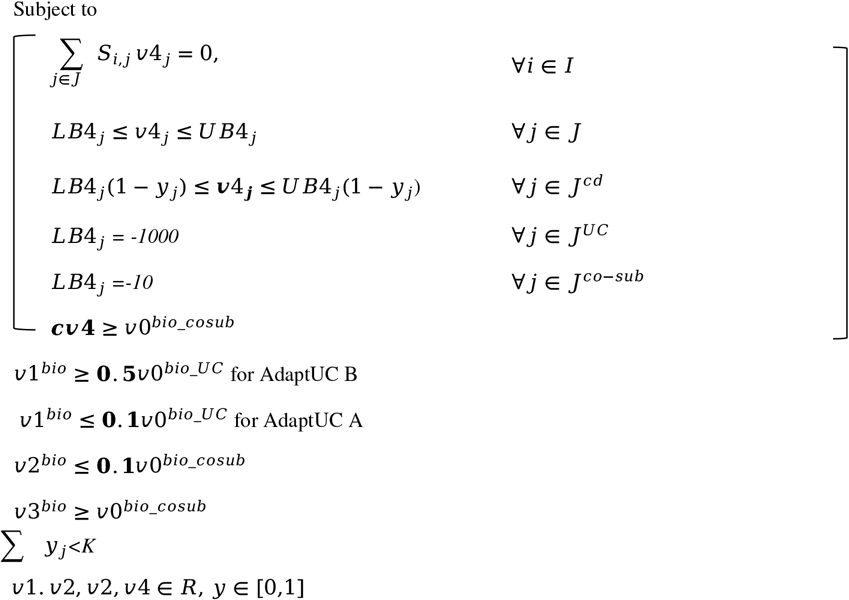

### Model Preprocessing

*E. Coli*: A methanol assimilation pathway consisting of three heterogeneous reactions encoded by RuMP methanol assimilation passway (*medh*: methanol dehydrogenase, *hps*:3-hexulose-6-phosphate synthase, and *phi*:3-hexulose-6-phosphate isomerase) were added to the *E. coli* model iML1515 ^21^. Two reactions, ‘PFL’ catalyzed by pyruvate formate lyase, and ‘OBTFL’ catalyzed by 2-oxobutanoate formate lyase,were deactivated in the model as they are only active under anaerobic condition. Additionally the reactions ‘DRPA’, ‘PAI2T’ were also removed since they are reported unrealistic ^22^ *C. glutamicum*: The genome scale metabolic model iCW773 ^23,24^ for *C. glutamicum* ATCC 13032 was utilized, methanol assimilation passways (*medh, hps*, and *phi*) were integrated to the model. To enable xylose utilization, a xylose transport reaction was also added.

For all models, non-essential carbon fixation reactions were deactivated except for those catalyzed by phosphoenolpyruvate carboxylase, carbamate kinase, isocitrate dehydrogenase, carbamoyl-phosphate synthase, pyruvate carboxylase, acetyl-CoA carboxylase, and methylmalonate-semialdehyde dehydrogenase.

### Evaluation metrics of the metabolic engineering strategies for the starting strains for ALE

AdaptUC offers different metabolic engineering strategies for specific hosts and substrates. However, selecting the most effective strategies for designing the initial strain for ALE requires careful assessment. For a successful co-substrate-aided ALE, the strain at the beginning of the ALE process must be able to grow on media containing the unadapted carbon source. Since it is unlikely for a starting strain to quickly uptake the unadapted carbon source, a low requirement for the unadapted source is crucial for survival. We quantify this with the Parsimonious Unadapted Carbon source and Co-substrate Assimilation Ratio (UC/Co), which is the minimum uptake rate of an unadapted carbon source relative to the minimum uptake rate of a co-substrate. A lower UC/Co indicates that less unadapted carbon source is required for the survival of the starting strain. To obtain UC/Co, we set a minimum specific growth rate of 0.1 hr^−1^ as a constraint for FBA to determine the minimum unadapted carbon source uptake rate. Subsequently, by fixing this minimum unadapted carbon source uptake rate, we determine the minimum co-substrate uptake rate. The ratio of these two uptake rates defines UC/Co.

Equally importantly, there must be sufficient evolutionary drive for the strain to utilize the unadapted substrate as the sole carbon source. This represents the selective pressure exerted on the microbial population to adapt and acquire traits necessary for utilizing previously unadapted carbon sources for growth. The response to selective pressure is described by the following equation:^25,26^

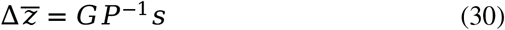

Where *s* is the selection differential (the growth rate difference between evolved and unevolved strains, ***Δ μ***) 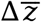 denotes the change of mean traits, (the change in uptake rate of the unadapted carbon source, ***Δ v***_***EX***_), and G and P are the matrices of genetic and phenotypic variance-covariance respectively^25,26^, The relative mean fitness gain from assimilating the unadapted carbon source can be expressed as:

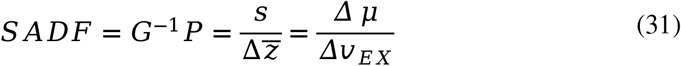

We define this relative mean fitness gain as Substrate Assimilation Driving Force (SADF). A higher SADF indicates greater selective pressure and a higher feasibility of evolution. To encompass the evolutionary driving force across various unadapted carbon source uptake rates, we compute the average SADF. This calculation involves determining the growth difference between the genetically engineered initial strain on a sole co-substrate (μ 2) and the evolved strains assumed to regain the growth rate achieved by the initial strain on a conventional carbon source (μ 0^*cosub*^), through assimilating the unadapted carbon source from a mixed medium containing both unadapted and conventional carbon sources. The denominator is the minimal uptake rate of the unadapted carbon source v^uc^. The average SADF can be formulated as:

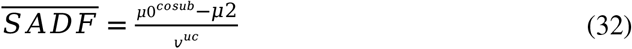

To calculate these growth rates and substrate uptake rates in eqn. (32), we employ FBA conducted on the genome-scale metabolic model. More detailed information can be found in the Method section.

## Acknowledgements

This work was supported by National Key Research and Development Program “Synthetic Biology” Project (2022YFA0912003) National Natural Science Foundation of China (32100035).

## Author contributions

**Jingyi Cai:** Conceptualization, Methodology, Software, Data curation, Visualization, Writing - original draft, Writing - review &Editing. **Jiayu Liu:** Data curation, Visualization. **Fan Wei:** Data curation, Visualization, **Wenjun Wu:** Software. **Wenqi Xu:** Visualization. **Zhitao Mao:** Writing - review &Editing **Qianqian Yuan:** Conceptualization, Visualization, Writing - review &Editing. **Hongwu Ma:** Conceptualization, Writing - review &Editing.

## Reference

1. Li, P., Bi, J., Liu, J., Zhu, Q., Chen, C., Sun, X., Zhang, J., and Han, B. (2022). In situ dual doping for constructing efficient CO2-to-methanol electrocatalysts. Nat Commun 13, 1–9. 10.1038/s41467-022-29698-3.

2. Ewis, D., Arsalan, M., Khaled, M., Pant, D., Ba-Abbad, M.M., Amhamed, A., and El-Naas, M.H. (2023). Electrochemical reduction of CO2 into formate/formic acid: A review of cell design and operation. Sep Purif Technol 316, 123811. 10.1016/j.seppur.2023.123811.

3. Boecker, S., Espinel-Ríos, S., Bettenbrock, K., and Klamt, S. (2022). Enabling anaerobic growth of Escherichia coli on glycerol in defined minimal medium using acetate as redox sink. Metab Eng 73, 50–57. 10.1016/j.ymben.2022.05.006.

4. Antonovsky, N., Gleizer, S., Noor, E., Zohar, Y., Herz, E., Barenholz, U., Zelcbuch, L., Amram, S., Wides, A., Tepper, N., et al. (2016). Sugar Synthesis from CO2 in Escherichia coli. Cell 166, 115–125. 10.1016/j.cell.2016.05.064.

5. Keller, P., Noor, E., Meyer, F., Reiter, M.A., Anastassov, S., Kiefer, P., and Vorholt, J.A. (2020). Methanol-dependent Escherichia coli strains with a complete ribulose monophosphate cycle. Nat Commun 11, 1–10. 10.1038/s41467-020-19235-5.

6. Tamari, Z., Yona, A.H., Pilpel, Y., and Barkai, N. (2016). Rapid evolutionary adaptation to growth on an “unfamiliar” carbon source. BMC Genomics 17, 1–7. 10.1186/s12864-016-3010-x.

7. Mao, Y., Li, G., Chang, Z., Tao, R., Cui, Z., Wang, Z., Tang, Y.J., Chen, T., and Zhao, X. (2018). Metabolic engineering of Corynebacterium glutamicum for efficient production of succinate from lignocellulosic hydrolysate. Biotechnol Biofuels 11, 1–17. 10.1186/s13068-018-1094-z.

8. Yu, H., and Liao, J.C. (2018). A modified serine cycle in Escherichia coli coverts methanol and CO2 to two-carbon compounds. Nat Commun 9, 1–24. 10.1038/s41467-018-06496-4.

9. Keller, P., Reiter, M.A., Kiefer, P., Gassler, T., Hemmerle, L., Christen, P., Noor, E., and Vorholt, J.A. (2022). Generation of an Escherichia coli strain growing on methanol via the ribulose monophosphate cycle. Nat Commun 13, 1–13. 10.1038/s41467-022-32744-9.

10. Chen, F.Y.H., Jung, H.W., Tsuei, C.Y., and Liao, J.C. (2020). Converting Escherichia coli to a Synthetic Methylotroph Growing Solely on Methanol. Cell 182, 933–946.e14. 10.1016/j.cell.2020.07.010.

11. Kim, S.J., Yoon, J., Im, D.K., Kim, Y.H., and Oh, M.K. (2019). Adaptively evolved Escherichia coli for improved ability of formate utilization as a carbon source in sugar-free conditions. Biotechnol Biofuels 12, 1–12. 10.1186/s13068-019-1547-z.

12. Kim, W., Lindner, S.N., Aslan, S., Yishai, O., Wenk, S., Schann, K., and Bar-Even, A. (2020). Growth of E. coli on formate and methanol via the reductive glycine pathway. Nat Chem Biol 16, 538–545. 10.1038/s41589-020-0473-5.

13. Wang, Y., Fan, L., Tuyishime, P., Liu, J., Zhang, K., Gao, N., Zhang, Z., Ni, X., Feng, J., Yuan, Q., et al. (2020). Adaptive laboratory evolution enhances methanol tolerance and conversion in engineered Corynebacterium glutamicum. Commun Biol 3. 10.1038/s42003-020-0954-9.

14. Orth, J.D., Thiele, I., and Palsson, B.O. (2010). What is flux balance analysis? Nat Biotechnol 28, 245–248. 10.1038/nbt.1614.

15. Mao, Z., Yuan, Q., Li, H., Hang, Y., Huang, Y., Yang, C., Wang, R., Yang, Y., Wu, Y., Yang, S., et al. (2023). CAVE: A cloud-based platform for analysis and visualization of metabolic pathways. Nucleic Acids Res 51, W70–W77. 10.1093/nar/gkad360.

16. Gawand, P., Hyland, P., Ekins, A., Martin, V.J.J., and Mahadevan, R. (2013). Novel approach to engineer strains for simultaneous sugar utilization. Metab Eng 20, 63–72. 10.1016/j.ymben.2013.08.003.

17. Bennett, R.K., Dillon, M., Gerald Har, J.R., Agee, A., von Hagel, B., Rohlhill, J., Antoniewicz, M.R., and Papoutsakis, E.T. (2020). Engineering Escherichia coli for methanol-dependent growth on glucose for metabolite production. Metab Eng 60, 45–55. 10.1016/j.ymben.2020.03.003.

18. Chen, C.T., Chen, F.Y.H., Bogorad, I.W., Wu, T.Y., Zhang, R., Lee, A.S., and Liao, J.C. (2018). Synthetic methanol auxotrophy of Escherichia coli for methanol-dependent growth and production. Metab Eng 49, 257–266. 10.1016/j.ymben.2018.08.010.

19. Meyer, F., Keller, P., Hartl, J., Gröninger, O.G., Kiefer, P., and Vorholt, J.A. (2018). Methanol-essential growth of Escherichia coli. Nat Commun 9. 10.1038/s41467-018-03937-y.

20. Cai, J., Liao, X., Mao, Y., Wang, R., Li, H., and Ma, H. (2023). Designing gene manipulation schedules for high throughput parallel construction of objective strains. Biotechnol J 18, 2200578. 10.1002/biot.202200578.

21. Monk, J.M., Lloyd, C.J., Brunk, E., Mih, N., Sastry, A., King, Z., Takeuchi, R., Nomura, W., Zhang, Z., Mori, H., et al. (2017). iML1515, a knowledgebase that computes Escherichia coli traits. Nat Biotechnol 35, 904–908. 10.1038/nbt.3956.

22. He, H., Höper, R., Dodenhöft, M., Marlière, P., and Bar-Even, A. (2020). An optimized methanol assimilation pathway relying on promiscuous formaldehyde-condensing aldolases in E. coli. Metab Eng 60, 1–13. 10.1016/j.ymben.2020.03.002.

23. Zhang, Y., Cai, J., Shang, X., Wang, B., Liu, S., Chai, X., Tan, T., Zhang, Y., and Wen, T. (2017). A new genome-scale metabolic model of Corynebacterium glutamicum and its application. Biotechnol Biofuels 10. 10.1186/s13068-017-0856-3.

24. Niu, J., Mao, Z., Mao, Y., Wu, K., Shi, Z., Yuan, Q., Cai, J., and Ma, H. (2022). Construction and Analysis of an Enzyme-Constrained Metabolic Model of Corynebacterium glutamicum. Biomolecules 12, 1499. 10.3390/biom12101499.

25. Lande, R., and Arnold, S.J. (1983). The Measurement of Selection on Correlated Characters. Evolution (N Y) 37, 1210. 10.2307/2408842.

26. Rausher, M.D. (1992). The measurement of selection on quantitative traits: biases due to environmental covariances between traits and fitness. Evolution (N Y) 46, 616–626. 10.1111/j.1558-5646.1992.tb02070.x.

