## Supplementary material for "Probing Biomass Precursor Synthesis as a Key Factor in Microbial Adaptation to Unadapted Carbon Sources": SI Notes

#### 1， Comparing literature-reported strategies with AdaptUC predictions

As shown in SI Table 1, the reactions that need to be knocked out in the strategies reported in the literature are not entirely consistent with those predicted by AdaptUC. Below, we will analyze the causes for these discrepancies.

SI Table 1, Comparison between literature reported and AdaptUC predicted metabolic modifications for construction of development of methylotrophic strains, “**^a”^** represent the strategy predicted by AdaptUC B, all others are predicted by AdaptUC A

| Organism | Co-substrate | Literature Reported Gene deletion | Reference | AdaptUC designed Gene deletions |
| --- | --- | --- | --- | --- |
| *E. coli* | Xylose | (RpiA b2914) ,(RpiB b4090) | Bennett, et al (2020)^1^ | (RpiA b2914) ,(RpiB b4090) |
|  | Ribose | (Rpe b3386) | Bennett, et al(2020)^1^ | (Rpe b3386) |
|  | Gluconate | (Edd b1851), (RpiA b2914) , (RpiB b4090),(Mdh b3236) | Meyer, et al(2018)^2^ | (EDD b1851), (RpiA b2914) ,(RpiB b4090) |
|  | Glucose | (Pgi b4025), (Edd b1851),(RpiA b2914), (RpiB b4090),(FrmA b0356) | Chen, et al(2018).^3^ | (Pgi b4025),(Edd b1851),(RpiA b2914), (RpiB b4090),HEX7(b0394) |
|  | Pyruvate | (FrmA b0356)，(Tpi b3919) | Keller, et al (2022)^4^ | (Tpi b3919),(GAPP b1727)**^a^** |
| C. *glutamicum* | Xylose | (RpiB, Cgl2423) (AdhE,Ald) | Wang, et al (2020)^5^ | (RPI Cgl2423 ) |

HEX7 (Hexokinase): In case 4, when using glucose as a co-substrate, the predicted strategy requires knocking out HEX7. However, this reaction has not been reported as necessary in experimental studies. Hexokinase phosphorylates hexose to produce fructose 6-phosphate (f6p). The gene for the HEX7 protein is identified as 'mak' in the iML1515 model, but no enzyme activity data such as kcat, MW, and Km for mak in Escherichia coli can be found in databases like UniProt, KEGG, and BRENDA. We discovered that the yajF gene in E. coli K12 corresponds biologically to the mak gene in iML1515. To further verify their similarity, we conducted a BLAST comparison on NCBI, which revealed a 100% similarity in their DNA sequences with an E-value of 0. This suggests that they are likely different names for the same gene, then we summarized *E. coli* sourced kinetic parameters of these two enzyme from Brenda, as show in SI Table 2

SI Table 2 enzyme activity of glk and YajF

| Enzyme | Substrates | *k_cat_* (s^-1^) | *K_m_*_,_ *_sugar_* (M) | *K_m_*_,_ *_ATP_* (M) | *k_cat_*/*K_m, sugar_* (M^-1^s^-1^) |
| --- | --- | --- | --- | --- | --- |
| glk | D-glucose | 410 | 7.6$\times$10^-5^ | 5.9$\times$10^-2^ | 5.4$\times$10^6^ |
| YajF | D-glucose | 12 | 5.9$\times$10^-2^ | 1.2$\times$10^-2^ | 2.0$\times$10^2^ |
|  | D-fructose | 14 | 1.3$\times$10^-3^ | 1.8$\times$10^-3^ | 1.1$\times$10^4^ |

under 25℃，pH= 7.6，0.3-0.5M Tris-HCl buffer

Based on SI Table 2, the kcat value of the glk enzyme in the PTS pathway is an order of magnitude higher than that of the YajF enzyme. The kcat/Km ratios for glucose and fructose are 4 to 2 orders of magnitude higher, respectively. This indicates that the Glk enzyme is relatively superior in catalytic activity, whereas the catalytic activity of the YajF enzyme is comparatively weaker. This result suggests that due to the very low abundance and catalytic activity of the native Mak (YajF) enzyme in E. coli, and the competition from GLK, there will be no significant metabolic flux through HEX7 in the actual cell without modification or overexpression. Therefore, even if HEX7 is not knocked out, it will not affect the overall metabolic pathway. The overprediction of HEX7 knockout is actually due to the stoichiometric model not accounting for reaction kinetics.

frmA （formaldehyde dehydrogenase）:

In cases 4, 5, and 6 of SI Table 1, the predicted results did not deactivate formaldehyde dehydrogenase (FrmA in E. coli and AdhE in C. glutamicum) as observed in experiments. Formaldehyde dehydrogenase is an essential enzyme in the formaldehyde detoxification pathway. This pathway detoxifies the cytotoxic formaldehyde, which spontaneously reacts with reduced glutathione to form S-hydroxymethylglutathione. Formaldehyde dehydrogenase (FrmA) then oxidizes S-hydroxymethylglutathione to S-formylglutathione. Finally, S-formylglutathione hydrolase hydrolyzes this compound to formate and reduced glutathione, completing the cycle.

This indicates that the formaldehyde detoxification process competes with the formaldehyde assimilation reaction (HPS-PHI). Upon reviewing the BRENDA database, we obtained the following data:：

SI Table 3 enzyme activity of hps and frmA

| BIGG ID/genes | Species | gene | *k_cat_*/*K_m_*  (m^–1^. s^–1^) |
| --- | --- | --- | --- |
| HPS-PHI^[88]^/hps | *Methylotuvimicrobium alcaliphilum* | 3-hexulose-6-phosphate synthase | 7.9 × 10^3^ |
| FALDH2^[89]^ (*frmA*) | *Escherichia coli* | Formaldehyde dehydrogenase | 1.65 × 10^6^ |

The kcat/Km value for the HPS-PHI reaction is three orders of magnitude lower than that for the FALDH2 (frmA) reaction, and 1-2 orders of magnitude lower than for SFGTHi. This indicates that the catalytic efficiency of Rump-related enzymes is significantly lower than that of E. coli's native formaldehyde detoxification system. Consequently, knocking out frmA facilitates methanol assimilation through the HPS-PHI pathway.

In the AdaptUC predictions, we found that only a minimal amount of metabolism occurred via the formaldehyde detoxification pathway (e.g., in the strategies shown in Fig. 3B-C, only 0.24% of formaldehyde is metabolized through frmA; see SI Fig. 1 for detailed metabolic flux). This indicates that the results predicted by AdaptUC do not actually conflict with the knockout of frmA. The absence of ΔfrmA in the predictions is due to the lack of kinetic information in the stoichiometric model, rather than a fundamental inconsistency with the need for frmA knockout.

GAPP（Glyceraldehyde-3-phosphate phosphatase）:

In case 5 of SI Table 1, manual inspection identified the presence of GAPP as the reason why the E. coli model could still grow on pyruvate after the Tpi reaction was knocked out. GAPP can substitute for Tpi to produce D-glyceraldehyde, which can further convert to fructose-6-phosphate, participate in the pentose phosphate pathway (PPP), and synthesize biomass precursors. This enables E. coli to grow on pyruvate alone, conflicting with the constraints of AdaptUC-B, resulting in the deactivation of GAPP. However, experimental results show that not actively knocking out GAPP does not enable Tpi-deficient E. coli to grow on pyruvate, casting doubt on the existence of the GAPP reaction in E. coli.

Upon investigation, GAPP is annotated in the BIGG database as gene b1727, which catalyzes the YniC enzyme and belongs to the HAD superfamily. However, a 2009 study by Kuznetsova^6^ on the enzymatic activity and catalytic mechanisms of YniC and the HAD family found no evidence of glyceraldehyde 3-phosphate in their substrate spectra. Similarly, the source literature for this reaction in the iML1515 model does not mention YniC catalyzing 3-phosphoglyceraldehyde^7^. Although the thermodynamic parameters theoretically allow this reaction to occur, a comprehensive review of the literature and GPR relationships casts doubt on the existence of the GAPP reaction in E. coli.

Therefore, AdaptUC-B's knockout of GAPP to correct the model is reasonable. This also highlights the need for strict verification of the existence of reactions in metabolic network models to ensure predictive accuracy.

**2, Lists of Seleted promising strategies predicted by AdaptUC for *E.coli* and *C. glutamicum***

SI Table 4. AdaptUC designed strategies with SADF greater than 5 for *E. coli*

| SADF | Algorithm | Reaction Deletion | Co-substrate |
| --- | --- | --- | --- |
| 24.896 | AdaptUC_A | FBP,RPE,F6PA | D-Gluconate |
|  |  | FBA,RPE,F6PA | D-Gluconate |
| 23.4945 | AdaptUC_A | FBP,TKT1,F6PA | D-Gluconate |
|  |  | FBA,TKT1,F6PA | D-Gluconate |
| 23.3672 | AdaptUC_A | RPE,TPI,GAPP | D-Gluconate |
| 22.5643 | AdaptUC_A | TKT1,TPI,GAPP | D-Gluconate |
|  |  | TKT1,ALCD19,TPI | D-Gluconate |
| 11.9555 | AdaptUC_B | FBA,GLYCDx,ALKP | Glycerol |
|  | AdaptUC_A | FBP,ALCD19,F6PA,G3PD2 | Glycerol |
|  |  | FBP,F6PA,ACONTa | L-Glutamate |
|  | AdaptUC_B | FBA,F6PA | Glycerol , Pyruvate , L-Glutamate , Succinate , Acetate |
|  |  | FBA,F6PA,GND | D-Gluconate |
|  | AdaptUC_A | PFK_3,FBA,EDD,F6PA | Glycerol |
|  | AdaptUC_B | FBA,GLYCDx,GND,ALKP | D-Gluconate |
|  |  | FBP,F6PA,GND | D-Gluconate |
|  |  | FBP,GLYCDx,ALKP | Glycerol |
|  |  | FBP,F6PA | Acetate , Glycerol , Pyruvate , Succinate , L-Glutamate |
| 10.2852 | AdaptUC_B | TPI,GAPP* | Succinate , Pyruvate* , L-Glutamate , |
|  |  | GND,TPI,GAPP | D-Gluconate |
|  |  | ALCD19,TPI | Pyruvate , L-Glutamate , Succinate |
|  |  | ALCD19,GND,TPI | D-Gluconate |

SI Table 5. AdaptUC designed strategies with SADF greater than 5 for *C. glutamicum*

| SADF | mode | delRxns | co_sub |
| --- | --- | --- | --- |
| 13.6903 | AdaptUC_A | **RPI** | Succinate , **Pyruvate** , Acetate , D-Glucose , Glycerol , L-Glutamate , D-Fructose |
| 9.8614 | AdaptUC_B | **FBP** | Succinate , Glycerol , L-Glutamate , Acetate , **Pyruvate** |
|  | AdaptUC_A | FBP,PFK | Succinate , Acetate , Glycerol , L-Glutamate , Pyruvate |
|  |  | FBP,PDH,POX | Acetate |
|  |  | FBA | Succinate , Glycerol , Acetate , L-Glutamate , Pyruvate |
|  |  | ACONTa,FBP | L-Glutamate |
| 9.2618 | AdaptUC_A | TPI,TRPS2 | Acetate |
|  |  | TPI | Succinate , L-Glutamate , Pyruvate |
| 8.9139 | AdaptUC_A | G3PD2,FBP | Glycerol |

**3, miscellaneous tables**

SI Table 6, Details of x-axis labels for Fig 6

| Strategy Groups | Strategies(Cosubstrate: Targets deleted) |
| --- | --- |
| group1 | xyl:TKT1;xyl:RPI;xyl:PFK_3,TALA;xyl:TALA,FBA3;rib:TKT1;rib:RPE;rib:TALA,FBA3;rib:PFK_3,TALA;glcn:EDA,TKT1;glcn:EDA,RPE;glcn:EDA,RPI;glcn:TKT1,EDD;glcn:EDD,RPI;glcn:RPE,EDD;glcn:PFK_3,TALA,EDD;glcn:TALA,EDD,FBA3 |
| group2 | xyl:ENO,PPS,TKT2 |
| group3 | glu:ICDHyr,PGM,ACONTa |
| group4 | ac:CPPPGO,MGSA,PGL,GAPD,FBA3,PDH |
| group5 | ac:MGSA,PGK,G6PDH2r;ac:MGSA,GAPD,G6PDH2r;ac:MGSA,EDA,GAPD;ac:MGSA,EDD,GAPD;ac:MGSA,PGK,EDA;ac:MGSA,PGK,EDD;pyr:MGSA,PGK,EDD;pyr:MGSA,PGK,EDA;pyr:MGSA,PGK,G6PDH2r;pyr:MGSA,GAPD,G6PDH2r;pyr:MGSA,PGK,PGL;pyr:MGSA,EDD,GAPD;succ:MGSA,PGK,PGL;succ:MGSA,PGK,EDA;succ:MGSA,PGK,G6PDH2r;succ:MGSA,PGL,GAPD;succ:MGSA,EDA,GAPD;succ:MGSA,PGK,EDD;succ:PPS,PGK,ICL;succ:MGSA,EDD,GAPD;glu:PGK,ACONTb;glu:GAPD,ACONTa;glu:GAPD,ACONTb;glu:PGK,ACONTa;glu:CS,PGK;glu:ICDHyr,GAPD;glu:ICDHyr,PGK;glu:CS,GAPD;glu:MGSA,PGK,EDA;glu:MGSA,EDA,GAPD;glu:MGSA,PGK,PGL;glu:MGSA,PGK,G6PDH2r;glu:MGSA,PGL,GAPD |
| group6 | glcn:TPI,RPI,GAPP |
| group7 | glyc:PFK_3,FBA,EDD,F6PA;glyc:FBA,EDD,F6PA,FBA3;glu:FBP,F6PA,ACONTb;glcn:FBP,F6PA,RPI |
| group8 | glcn:FBA,TKT1,F6PA;glcn:FBP,TKT1,F6PA |
| group9 | glcn:FBA,RPE,F6PA;glcn:FBP,RPE,F6PA |

SI Table 7, Details of y-axis labels for Fig 6

| Precursor Groups | Biomass Precursors |
| --- | --- |
| precursor group 1 | btn;ribflv;succoa;amet;thf;mlthf;10fthf;fad;sheme;nadp;coa;nad;dttp;dctp;datp;dgtp;his;ctp;utp;gtp |
| precursor group 2 | pro;thr;asp;asn;ile;glu;gln;ala;val;lys;arg;leu |

**Reference**

1.	Bennett, R. K. *et al.* Engineering Escherichia coli for methanol-dependent growth on glucose for metabolite production. *Metab Eng* **60**, 45–55 (2020).

2.	Meyer, F. *et al.* Methanol-essential growth of Escherichia coli. *Nat Commun* **9**, (2018).

3.	Chen, C. T. *et al.* Synthetic methanol auxotrophy of Escherichia coli for methanol-dependent growth and production. *Metab Eng* **49**, 257–266 (2018).

4.	Keller, P. *et al.* Generation of an Escherichia coli strain growing on methanol via the ribulose monophosphate cycle. *Nat Commun* **13**, 1–13 (2022).

5.	Wang, Y. *et al.* Adaptive laboratory evolution enhances methanol tolerance and conversion in engineered Corynebacterium glutamicum. *Commun Biol* **3**, (2020).

6.	Kuznetsova, E. Activity-based Functional Annotation of Unknown Proteins: HAD-like hydrolases from E. coli and S. cerevisiae. (2009).

7.	Kuznetsova, E. *et al.* Genome-wide Analysis of Substrate Specificities of the Escherichia coli Haloacid Dehalogenase-like Phosphatase Family. *Journal of Biological Chemistry* **281**, 36149–36161 (2006).
